# Analysis of cancer mutations introduced into the Drosophila Notch Negative Regulatory Region uncovers a diversity of regulatory outcomes

**DOI:** 10.1101/2025.09.01.673436

**Authors:** Hideyuki Shimizu, Martin Baron

## Abstract

Activating mutations of Notch are drivers of the blood cell cancer, T-ALL, and some solid tumours. The Negative Regulatory Region (NRR) of the extracellular domain (ECD) and the PEST region of the intracellular domain (ICD) are mutation hot spots which can act synergistically in T-ALL. The NRR, comprised of a Heterodimerisation Domain (HD) and three Lin12/Notch repeats (LNR A-C), masks the S2 cleavage site, normally only exposed following ligand binding and cleaved as the first step that ultimately leads to ICD release. *Drosophila* mutants have played a key role in analysing Notch structure/function but there have been few mutational studies of the NRR. Here we expressed, in S2 cells, over 20 cancer mutations located in the HD, LNR and LNR/HD interface, introduced into *Drosophila* Notch. Mutations in the HD domain core did not activate, likely due to absence, in *Drosophila*, of an S1 cleavage within the HD, required for mammalian Notch activity. In contrast, mutations in the LNR/HD interface behaved similarly to T-ALL, activating constitutively with no further ligand-induction and were synergistic with PEST deletion. Mutations of surfaced-exposed residues of LNR-C, also activated constitutively but remained inducible both by ligand and by an intracellular endocytic regulator, Deltex, and were not synergistic with PEST deletions. These mutations caused elevated Notch levels and decreased turnover, suggesting a novel regulatory mechanism. Our results therefore uncover a variety of outcomes arising from perturbations of the NRR and will facilitate the establishment of *Drosophila* cancer models and the development of mutant-specific approaches to effective therapies.

## Introduction

Notch proteins are single-pass transmembrane receptors that can be activated by either transmembrane ligands such as Delta (Dl) and Serrate/Jagged, or by endocytosis in a ligand-independent manner (Sachan et al., 2024; Zhou et al., 2022). In the mechanotransduction model of Notch signalling, force-dependent activation machinery ensures precise regulation of the signal in response to specific ligand-receptor interactions (Sprinzak and Blacklow, 2021; Suarez Rodriguez et al., 2023). The ligand binding meditates an endocytic pulling force on the Notch receptor, which is detected by the mechanosensitive Negative Regulatory Region (NRR) which comprises of three Lin12/Notch repeats (LNR-A, LNR-B and LNR-C) together with the heterodimerisation domain (HD). The latter is a domain which is cleaved at the S1 site by Furin during transit of Notch proteins through the endoplasmic reticulum, a step which is thought to be a prerequisite for efficient transport of vertebrate Notch proteins to the cell membrane (Logeat et al., 1998). However, this step appears not to be required for *Drosophila* Notch (Kidd and Lieber, 2002). The ligand-induced mechanical force triggers a conformational change that exposes the S2-cleavage site in the HD (Chen and Zolkiewska, 2011; Gordon et al., 2007; Gordon et al., 2015; Stephenson and Avis, 2012). After exposure of the S2 site, ADAM family proteases catalyse S2-cleavage (Brou et al., 2000), and this is subsequently followed by γ-secretase-dependent S3 cleavage (De Strooper et al., 1999). The latter releases the Notch Intracellular Domain (NICD) from the membrane, allowing it to translocate into the nucleus (Struhl and Adachi, 1998), where it acts as a transcriptional co-activator for target genes. The PEST domain at the C-terminal end of Notch plays a crucial role in regulating NICD stability and turnover by serving as a signal for ubiquitination and subsequent proteasomal degradation (Gupta-Rossi et al., 2001; Hubbard et al., 1997; Oberg et al., 2001). In addition to the above mechanism, two ligand-independent activation mechanisms have been identified which arise in different endocytic routes. Basal activation occurs on the endosome membrane following clathrin-independent endocytosis. Interaction of Notch with the cytoplasmic RING domain ubiquitin ligase, Deltex (Dx), stimulates ligand-independent activation by diverting Notch into a clathrin-dependent endocytic pathway and promoting Notch accumulation in a clathrin-rich endosomal membrane domain. It is retained in this membrane domain until the Notch extracellular domain (ECD) is removed following lysosomal fusion and the ICD is released by gamma-secretase cleavage (Shimizu et al., 2024; Shimizu et al., 2014).

T-cell acute lymphoblastic leukaemia (T-ALL) is an aggressive haematological cancer (Belver and Ferrando, 2016; Cordo et al., 2021). It originates from the malignant transformation of T-cell progenitors in the thymus and involves the rapid proliferation of immature T-cells in the bone marrow and blood. Despite remarkable progress in understanding the underlying molecular mechanisms and advances in therapeutic treatments, T-ALL still has a relatively poor prognosis. Notch signalling plays critical roles in T-cell and B-cell development in human adult haematopoiesis. NOTCH1 has a pivotal function in T-cell differentiation (Ng et al., 2021), and consistent with this, over 50% of T-ALL cases have been reported to have mutations or chromosomal rearrangements in the NOTCH1 gene locus (Breit et al., 2006; Weng et al., 2004). These mutations clearly cluster in the HD and the PEST domain regions and disruption of their functions thus activates Notch Signalling. In the NRR, the majority of these mutations are located in the hydrophobic core of the HD but cancer mutations in the LNRs have also been reported, especially in solid tumours (Table 1).

**Table 1.** List of human NOTCH1 cancer mutations and their *Drosophila* equivalents used in this study. Summary of 22 cancer-associated mutations analysed in this study. *Drosophila* Notch mutations were designed based on reported human NOTCH1 cancer mutations. Both conserved and non-conserved residues were included to examine how structural perturbations at equivalent positions affect signalling activity, independent of sequence conservation. The table lists each *Drosophila* mutation alongside its corresponding human mutation, structural location, associated cancer types, and relevant references.

| Drosophila | human NOTCH1 | location | cancer types | reference |
| --- | --- | --- | --- | --- |
| E1489V | E1456V | LNR-A | prostate cancer (PCa) | (Mateo et al., 2020) |
| D1497N | S1464N | LNR-A | PCa, squamous cell carcinoma (SCC) | (Mateo et al., 2020; South et al., 2014) |
| A1504V | A1471V | LNR-A | ampullary adenocarcinoma | (Gingras et al., 2016) |
| G1515K | N1482K | LNR-A/B | metastatic colorectal cancer (mCRC) | (Yaeger et al., 2018) |
| T1523M | T1491M | LNR-B | Merkel cell carcinoma | (Starrett et al., 2020) |
| N1529E | K1498E | LNR-B | skin melanoma | (Bi et al., 2017) |
| E1538R | S1507R | LNR-B | melanoma | (Abou Alaiwi et al., 2020) |
| D1548N | D1517N | LNR-B | head and neck (HN) SCC, T-cell acute lymphoblastic leukaemia (T-ALL) | (Neumann et al., 2015; Stransky et al., 2011) |
| E1553P | Q1522P | LNR-B/C | SCC | (South et al., 2014) |
| R1554C | R1523C | LNR-B/C | cervical SCC | (Zhang et al., 2016) |
| L1562P | L1531P | LNR-C | oral (O) SCC | (Izumchenko et al., 2015) |
| H1570P | H1539P | LNR-C | OSCC | (Izumchenko et al., 2015) |
| D1573V | D1542V | LNR-C | oesophageal SCC | (Zhang et al., 2015) |
| D1577G (414) | D1546G | LNR-C | T-ALL | (Asnafi et al., 2009) |
| Y1578R | Q1547R | LNR-C | mantle cell lymphoma | (Zhang et al., 2014) |
| F1617P | L1585P | HD-N | T-ALL | (Asnafi et al., 2009; Neumann et al., 2015; Weng et al., 2004) |
| R1626Q | R1594Q | HD-N | T-ALL, SCC, stomach | (Durinck et al., 2011; Ito et al., 2019; Zhang et al., 2016) |
| H1630P | R1598P | HD-N | T-ALL | (Asnafi et al., 2009; Neumann et al., 2015; Weng et al., 2004) |
| L1632P | L1600P | HD-N | T-ALL, breast | (Asnafi et al., 2009; Neumann et al., 2015; Pop et al., 2018; Weng et al., 2004) |
| R1633P | H1601P | HD-N | T-ALL | (Matteucci et al., 2010) |
| L1686P | L1678P | HD-C | T-ALL, mCRC, stomach | (Asnafi et al., 2009; Liu et al., 2014; Neumann et al., 2015; Weng et al., 2004; Yaeger et al., 2018) |
| E1705P | A1701P | HD-C | T-ALL | (Neumann et al., 2015; Weng et al., 2004) |

*Drosophila melanogaster* has contributed to groundbreaking discoveries in genetics, developmental biology, and disease mechanisms over the past century. The high degree of conservation between human and *Drosophila* Notch enable the latter to be used for probing structure/function relationships of Notch through mutational analysis (Nurmahdi et al., 2022; Yamamoto, 2020). However few mutations in the NRR region have been reported. Here we report a mutational analysis of the *Drosophila* Notch NRR in the S2 cell line, aiming to identify Notch mutants that can activate in a similar manner to human cancers. Using this *Drosophila* cell line, enables the consequences of the NRR mutations on basal, Dx-induced and ligand-induced signalling to be investigated separately (Shimizu et al., 2014), providing a means to distinguish different classes of outcomes. We introduced twenty-two mutations into *Drosophila* Notch that are equivalent to human NOTCH1 cancer mutations within the NRR. We tested outcomes on basal, constitutive activation and on Dx and Dl-inducibility. We found that mutations located in different regions of the NRR produced different responses in these assays. Whereas T-ALL mutations located inside the hydrophobic core of the HD exhibited no increase in signalling, cancer mutations lying between the LNR and HD showed high ligand-independent (basal) activation which lacked further inducibility. Interestingly, we uncover a novel regulatory region in LNR-C, where surface mutations increased constitutive basal signalling but remained highly inducible by both Dl and Dx. We found that these mutants most likely act by a novel regulatory mechanism that stabilises the Notch protein. These findings will contribute to establishing a T-ALL model in *Drosophila* and uncovering mutant-specific mechanisms will inform the development of targeted therapies.

## Results

### Analysis of ligand-independent Notch signalling by NRE-luciferase in S2 cells

The NRR consists of three LNR domains and the HD (heterodimerisation domain), which are highly similar between *Drosophila* Notch and human NOTCH proteins. Consequently, the signal activation mechanism is believed to be well conserved between these species. However, there is a notable difference in Notch protein maturation: while S1 cleavage by Furin-like convertases is essential for the maturation of mammalian Notch proteins (Logeat et al., 1998), the S1 processing of *Drosophila* Notch protein appears not to be essential (Kidd and Lieber, 2002). This difference suggests that HD mutations might have different effects on Notch signal activation in flies and humans.

We first tested whether the NRE-luciferase assay in the S2 cell line (Bray et al., 2005) could monitor the overactivation of existing *Drosophila* Notch gain-of-function alleles. Three mutations within the *Drosophila* Notch NRR—l1N-B (G1572V) (Lyman and Young, 1993), 414 (D1577G) (Brennan et al., 1997), and CC-SS (Lieber et al., 1993)—have been previously reported to exhibit gain-of-function phenotypes *in vivo* (Figure 1A). L1NB is located in LNR-C at the interface with the HD domain, while 414 is located within LNR-C forming part of calcium binding pocket (Gordon et al., 2009). An equivalent mutation in human Notch1 (D1546G) has been identified in T-ALL (Table 1). CC-SS is a double mutant that replaces the conserved disulphide bridge within the HD domain. In addition, a triple mutation LGI-AAA was generated to disrupt the gate keeper site (Gordon et al., 2007) which is located between the LNR-A and LNR-B domains and which normally shields the S2 site from ADAM protease-dependent cleavage (Figure 1A). As expected, all these mutants significantly increased basal Notch signalling (Figure 1B), even in the absence of the Notch ligand Dl, or Dx, confirming that the NRE reporter is effective in detecting Notch overactivation caused by NRR mutations. All four mutants displayed little further induction when exposed to Dl or Dx (Supplementary Figure S1A,B). The constitutive activity of L1N-B, 414 and CC-SS remained higher than even the Dl-induced WT signal. In contrast, CC-SS and LGI-AAA were not significantly higher than WT induced by Dx (Figure 1E). Interestingly, l1N-B and 414 activities, in the presence of Dx, were significantly less than Dx-induced WT (Figure 1E) and, in the case of 414, the expression of Dx significantly reduced Notch signalling compared to its own basal activity (Supplementary Figure S1 C).

**Figure 1.**
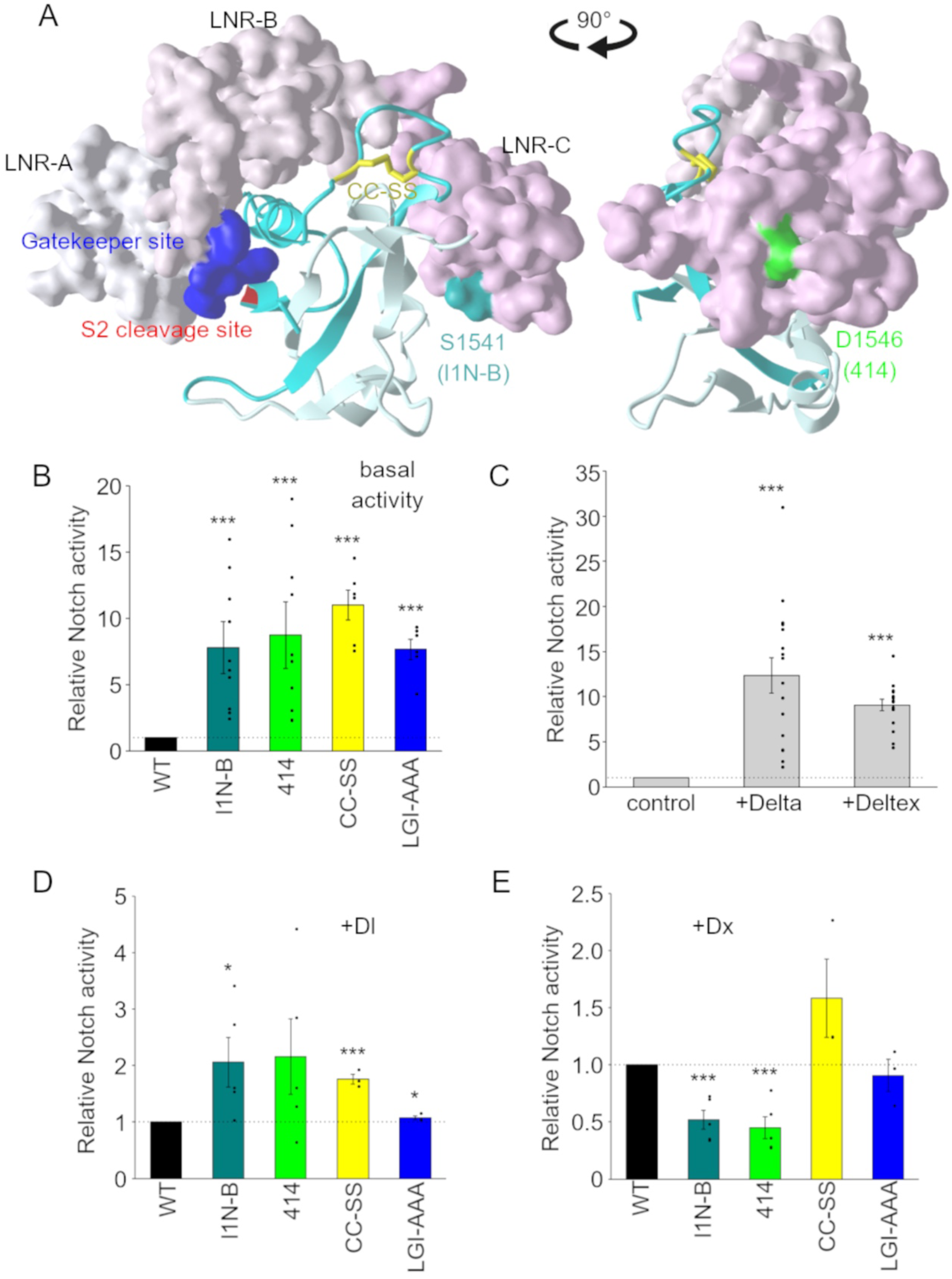
Ligand-independent activation of Notch signal by gain-of-function mutations in the *Drosophila* Notch NRR. **(A)** Front and side views of the human NOTCH1 NRR crystal structure, highlighting residues corresponding to four *Drosophila* gain-of-function mutations: l1N-B (G1572V; dark cyan), 414 (D1577G; green), CC-SS (C1693S and C1696S; yellow), and LGI>AAA (LGI1514–1516AAA; blue). The HD domains (HD-N; pale cyan and HD-C; cyan) are shown as ribbons to illustrate secondary structure and the positions of mutations within the fold, particularly those at the HD-N/HD-C interface. The LNR repeats are shown in space-filling representation to highlight their surface-exposed positions and facilitate visual distinction from the HD domains. All mutations in this study are uniquely colour-coded for consistency across structural representations and quantitative analyses, including NRE-luciferase assays and protein-level measurements. **(B)** Enhanced ligand-independent signalling by Notch gain-of-function mutants. NRE-luciferase assay in S2 cells show that all four gain-of-function Notch mutants exhibit significantly increased basal signalling activity compared to wild-type. **(C-E)** Ligand (Dl) and Dx-induced *Drosophila* Notch signal. Wild-type Notch signal can be further activated by Dl and Dx **(C)**. Dl- **(D)** and Dx- **(E)** induced Notch signal intensity of the four gain-of-function mutants were normalised by the signal from wild-type Notch. P < 0.05 (*), P < 0.01 (**), and P < 0.001 (***) by two tailed t-test, error bars are SEM.

This data indicates that the function of the LNR and gatekeeper site in shielding the S2 cleavage site is conserved between human and *Drosophila* Notch, highlighting an evolutionarily shared mechanism by which the NRR regulates Notch activation. The lack of inducibility to ligand or Dx suggests that the S2-cleavage site in the mutants is already exposed to ADAM10/Kuzbanian, mimicking the Notch activation mechanism triggered by its ligands. Thus, we concluded that *Drosophila* Notch and the NRE-luciferase assay provide a comprehensive platform to analyse human Notch cancer mutations in the NRR. The data also indicated some differences between mutant outcomes in response to regulators can be useful to classify the different cancer mutants.

### T-ALL mutations in the HD

Next, human NOTCH1 cancer mutations within the NRR were introduced into *Drosophila* Notch. In T-ALL patients, certain amino acids in the HD of NOTCH1 are frequently mutated. To study these mutations, we introduced seven of these hotspot mutations into equivalent positions within the HD of *Drosophila* Notch (F1617P, R1626Q, H1630P, L1632P, R1633P, L1686P, and E1705P) (Table 1, Supplementary figure S2, and Figure 2A). We examined the three different types of Notch signal activation pathways—basal, ligand-induced, and Dx-mediated endosomal activation—across all the Notch mutants. Surprisingly, mutations within the hydrophobic core of the HD (F1617P, L1632P, and L1686P) did not lead to an increase of the constitutive basal signal, which remained as WT or slightly reduced (Figure 2B). Two HD mutants H1630P and R1633P mutants are distinctive in their localisation because their side chains are surface-exposed, but these mutants also showed no increase signalling. In the presence of either Dl or Dx, all the above HD mutants signalled at only a small fraction of the induced WT level (Figure 2C,D), as there was little or no induction over basal levels (Supplementary figure S1D,E). In contrast, equivalent human NOTCH1 mutations are known to exhibit higher basal activity. This discrepancy is likely due to the different requirements for Notch S1 cleavage in *Drosophila* (Kidd and Lieber, 2002) compared to humans (Logeat et al., 1998). Notably, L1632P and L1686P showed partial accumulation in the ER and Golgi compared to WT, or another mutant, G1515K, located outside of the HD (Figure 2E). This suggests that quality control mechanisms (Phillips and Miller, 2020; Sun and Brodsky, 2019) might contribute to the lack of activation observed in these HD mutants. Interestingly, two mutants, R1626Q and E1705P, displayed much higher basal activity than WT. The side chains of the equivalent amino acids in NOTCH1 face the LNR domain, suggesting that they contribute to the interaction between the LNR and HD, whereas most of the other HD mutations are believed to be involved in stabilizing the interaction between HD-N and HD-C. Neither of these mutants responded to ligand or Dx-induction to increase their activity above their already high basal level (Supplementary figure S1D,E), suggesting that the S2 site was fully exposed and could not be further induced. Interestingly, in signal-induced conditions WT Notch activated at a higher level than both the R1626Q and E1705P (Figure 2C,D,) indicating cellular context is important in determining whether a particular mutation signals more, or less, than WT.

**Figure 2.**
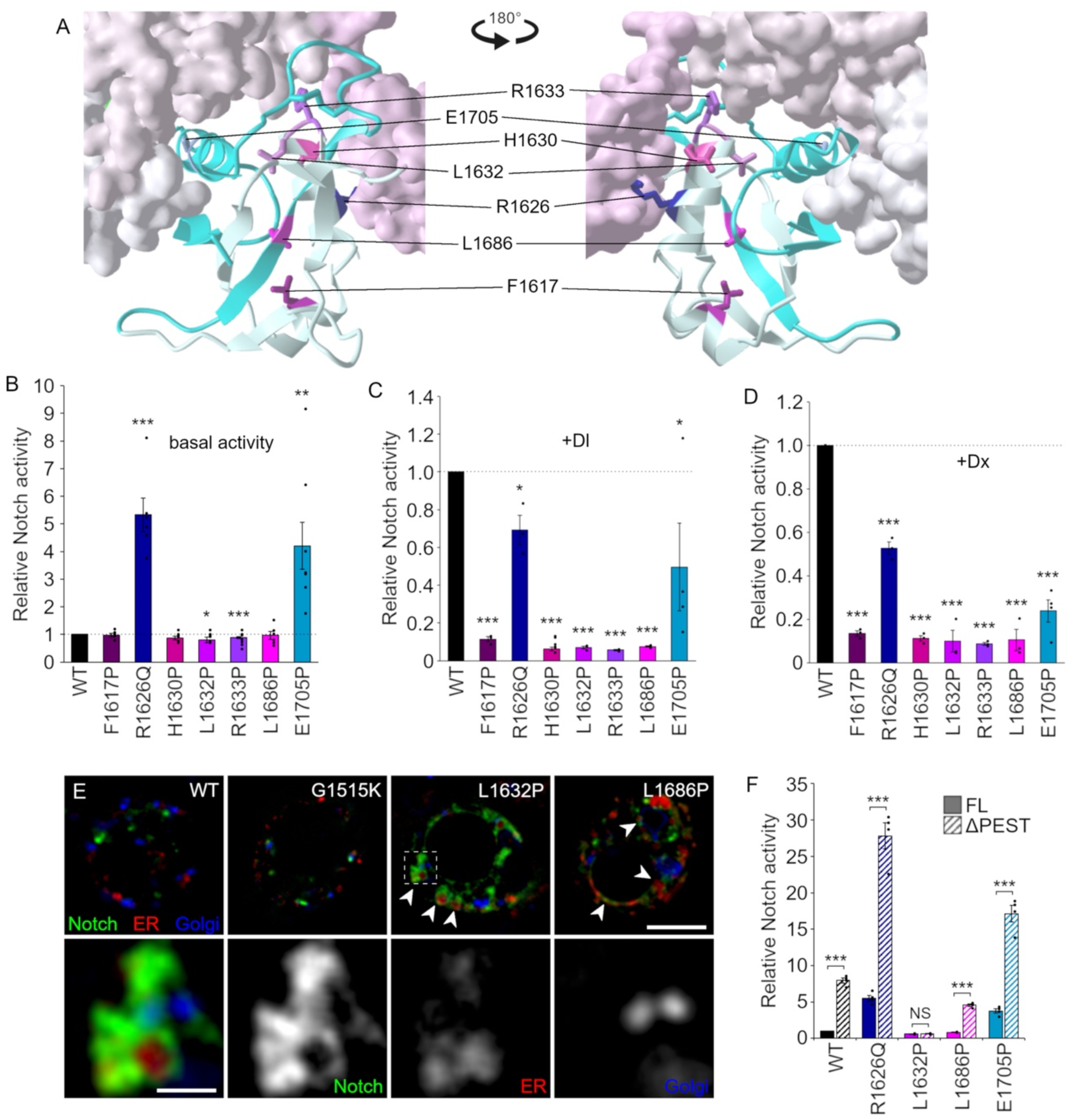
Functional analysis of T-ALL–derived heterodimerisation domain mutations in *Drosophila* Notch. **(A)** Positions of residues corresponding to seven *Drosophila* mutations (F1617P; dark magenta, R1626Q; dark blue, H1630P; strong pink, L1632P; soft magenta, R1633P; violet, L1686P; magenta, and E1705P; dark turquoise) in HD domain shown on human NOTCH1 NRR crystal structure. The selected residues represent recurrent, independently reported cancer-associated NOTCH1 HD mutations. F1617P, L1632P, and L1686P lie in the hydrophobic core; H1630P and R1633P are surface-exposed; and R1626Q, and E1705P are positioned at the LNR–HD boundary. **(B)** Basal Notch activity of the seven HD domain mutants, demonstrating that the only two mutants R1626Q and E1705P exhibit accelerated signal intensity in ligand-independent manner, like T-ALL mutants. **(C,D)** Notch signal activation of HD domain mutants by ligand **(C)** and Dx **(D)**, normalised by wild-type Notch activated by ligand and Dx, respectively. **(E)** Impaired secretory trafficking of Notch HD domain mutants due to ER/Golgi-retention. The localisations of two representative non-signalling HD mutants in conserved residues were investigated (L1632P, L1686P) and compared with the active signalling mutant G1515K. Upper panels: EGFP-tagged wild-type Notch and two HD domain mutants (L1632P and L1686P) were expressed in S2 cells and Notch protein localisation (EGFP, green) was analysed by immunofluorescence. The cells were stained with anti-PDI (for ER, red) and anti-GM130 (for Golgi, blue) antibodies. There was no clear difference between WT and G1515K localisation despite the intense basal signal of G1515K (Figure3B). Lower panels: higher magnification of the box shown in L1632P image. Scale bar: 5µm (upper panels) and 1µm (lower panels). **(F)** Effect of PEST domain truncation to ligand-independent Notch signal of HD domain mutants. PEST domain in the C-terminal region of Notch was removed from four representative mutants (R1626Q, L1632P, L1686P, and E1705P), and the basal signal was analysed by NRE-luciferase assay. Only R1626Q and E1705P show T-ALL mutants-like synergistic increase of the signal. P < 0.05 (*), P < 0.01 (**), and P < 0.001 (***) by two tailed t-test, error bars are SEM.

The PEST domain at the C-terminus of Notch serves as another negative regulatory region, facilitating the degradation of the Notch ICD fragment through a proteasome-dependent mechanism (Gupta-Rossi et al., 2001; Hubbard et al., 1997; Oberg et al., 2001). In T-ALL patients, HD mutations and PEST truncations are often found in cis, leading to more aggressive and treatment-resistant symptoms. Also, synergistic upregulation of ligand-independent NOTCH1 signalling has been observed with double mutants in cell culture assays (Sulis et al., 2008; Weng et al., 2004). To further analyse whether the *Drosophila* HD mutations can exhibit a similar synergy, a PEST domain truncation was combined with WT, R1626Q, L1632P, L1686P, and E1705P. Deletion of PEST region alone already stimulated the basal activity of WT Notch, however the NRE-luciferase assay revealed a clear synergy in Notch activity levels between removal of the PEST and the LNR-HD interaction mutations (R1626Q and E1705P). The hydrophobic core mutant L1632P showed no increase on removal of the PEST, while an increase in basal activity of L1686P appeared additive rather than synergistic (Figure 2F). Therefore, we concluded that two mutants (R1626Q and E1705P) could be used as T-ALL models in *Drosophila*, serving as alternatives to the typical T-ALL mutations in human NOTCH1, such as L1601P and L1678P.

### Cancer mutations in the LNR

Having characterised Notch HD mutations, we next analysed mutations located in the LNRs. Fourteen mutations identified in T-ALL and/or a range of solid tumours were selected from the NOTCH1 LNR-A to LNR-C regions (Table 1, Supplementary figure S2 and Figure 3A). These were located predominantly in surface-exposed regions of the LNR domains. The equivalent mutations were introduced into fly Notch and changes in the signal activation of each mutant were characterised using the NRE-luciferase assays for basal, Dx and Dl signalling modes. Among these LNR mutants, E1489V, D1497N, G1515K, E1553P, R1554C, L1562P, H1570P, and D1573V showed a higher basal Notch signal compared to the wild-type, while A1504V, N1529E, E1538R, and Y1578R had significantly lower basal signals (Figure 3B). Of the activating mutants, G1515K and E1553P displayed markedly stronger basal activation. The relative Notch activity in the presence of Dl and Dx of the mutant compared to WT also varied between different mutants, suggesting potential diversity in the final output of Notch signal intensity in individual tissues, depending on which pathways are predominantly used for their functions in different contexts (Figure 3C and 3D). Despite showing little further induction by Dl or Dx (Supplemental Figure S1 D,E), the E1553P mutant still had significantly higher activity than WT Notch when the latter was induced by Dl and both G1515K and E1553P mutants signalled stronger than WT when induced intracellularly by Dx (Figure 3C,D). Other mutants with higher basal activity than WT, i.e. E1489V, D1497N, R1554C, L1562P, showed varying amounts of inducibility, which resulted in them also having higher activity than WT for Dx and/or Dl-induced conditions (Figure 3C,D, Supplemental Figure S1 D,E)). Of the mutants that had lower basal activity than WT, the A1504, N1529E and Y1578R mutants were at least as responsive to ligand and Dx induction as was WT (Supplemental Figure S1), resulting in similar levels of activity to WT overall in these conditions. However, E1538R was unresponsive to either Dl or Dx and its activity was therefore significantly below the ligand/Dx-induced level of activity of WT Notch. The remaining LNR mutants T1523M, D1548N which showed no difference to WT for basal signalling levels had very different outcomes in signal-inducing conditions. While T1523M was similarly activated by Dx and Dl compared to WT, the D1548N mutant was not inducible by either Dx or Dl (Figure 3C,D, Supplemental Figure S1).

**Figure 3.**
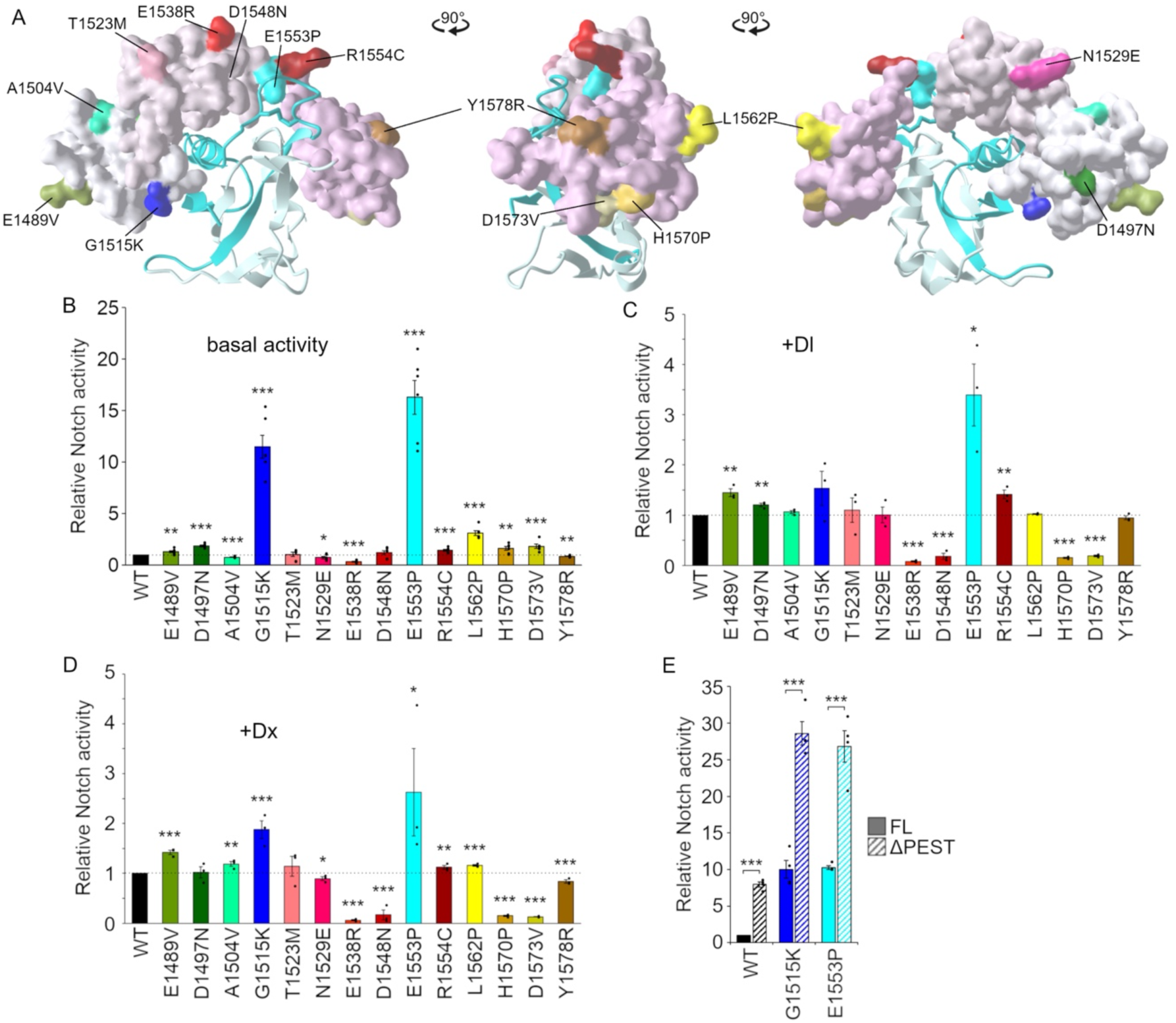
Structure-function analysis of the *Drosophila* Notch LNR domain via cancer-derived mutational mapping. **(A)** Distribution of cancer-associated LNR mutations on the human NOTCH1 NRR Structure. Fourteen mutations (E1489V; olive, D1497N; dark green, A1504V; aquamarine, G1515K; blue, T1523M; salmon, N1529E; red pink, E1538R; crimson, D1548N; maroon, E1553P; cyan, R1554C; brown, L1562P; yellow, H1570P; strong yellow, D1573V; dark yellow, and Y1578R; tan) were introduced into *Drosophila* Notch, based on cancer-associated variants reported in the human NOTCH1 LNR domain. The D1548N mutation, which contributes to a calcium binding site, is buried within the LNR and is not visible in the figure. **(B)** Basal Notch activity of the LNR domain mutants, demonstrating that the only two mutants G1515K and E1553P exhibit apparently high signal intensity in ligand-independent manner. **(C,D)** Notch signal activation of the LNR mutants by ligand **(C)** and Dx **(D)**, normalised by wild-type Notch activated by ligand and Dx, respectively. **(E)** Synergistic enhancement of ligand-independent Notch signalling by combined LNR mutations and PEST domain truncation. The effect of PEST domain truncation on the basal activity of two gain-of-function LNR mutants (G1515K and E1553P) was assessed using the NRE-luciferase assay. The combination led to a synergistic increase in ligand-independent signalling, resembling the behaviour of T-ALL–associated NOTCH1 mutations. P < 0.05 (*), P < 0.01 (**), and P < 0.001 (***) by two tailed t-test, error bars are SEM.

The side chains of G1515K and E1553P, which have the highest basal signals (Figure 3B), as well as G1572V (l1N-B allele, Figure 1), are oriented toward the HD, indicating that these residues may contribute to the protection of the S2-cleavage site through LNR-HD interaction, similar to R1626Q and E1705P in the HD. These mutations also showed synergistic increase of basal activity when combined with PEST domain truncation (Figure 3E), demonstrating that LNR-HD interaction mutants behave like typical T-ALL mutants.

### LNR-C Interface mutants

We found it interesting that certain LNR-C mutations have been reported to activate human Notch but are not directly involved in the LNR-HD interaction. Previous crystal structure analyses of human NOTCH1-3 NRRs have shown that some residues which cause activation when mutated are located on the lateral surface of LNR-C (Gordon et al., 2009; Xu et al., 2015) (Figure 3A). Intriguingly, these analyses have revealed that the LNR-C domains typically have a surface area that mediates homodimerisation between two NRRs, at least in the context of the crystal structures, with H1539 in NOTCH1 (equivalent to H1570 in *Drosophila* Notch) playing a key role. This interface is highly conserved across NOTCH1-3 and various species, including *Drosophila*. For instance, in NOTCH1, two main residues (Y1535 and H1539) and an auxiliary residue (Y1532) are predicted to be crucial for Notch homodimerisation. This published data therefore suggested that these LNR-C mutants may impact Notch signalling through an unknown mechanism involving Notch-Notch interaction.

To explore the negative regulatory roles of the LNR-C further we generated *Drosophila* Notch mutants in putative interface residues equivalent to Y1532, Y1535, and H1539 (i.e. F1563A, Y1566A, and H1570A) (Figure 4A), note that a H1539P mutation has been identified in T-ALL (Table1, Figure 3A). Y1532 and Y1535 are not mutated in human cancers and therefore could not be assessed through patient-derived variants. Alanine substitution provides a controlled way to probe their contribution to NRR integrity and activation sensitivity by selectively removing their side-chain interactions while preserving overall fold. All three mutants exhibited high basal Notch signal activity in the NRE-luciferase assay (Figure 4B), consistent with findings from a previous mammalian Notch study (Xu et al., 2015). Interestingly, unlike most of the gain-of-function NRR mutants, both F1563A and H1570A mutants showed similar levels of inducibility by Dl and Dx as the WT (Supplemental Figure S1F,G)), despite having an already high basal level of activity. Hence, their signalling levels in the presence of these stimuli were substantially higher than WT (Figure 4 C,D). The Y1566A mutant retained around 25% of WT inducibility by Dl but was barely responsive to Dx (Supplemental Figure S1). However, because of its already high basal levels, the activity of Y1566A was still significantly higher than WT in both these signal-inducing conditions. (Figure 4C,4D). These results suggested a new mechanism, distinct from S2-cleavage site exposure or HD destabilization, might be involved in driving the higher levels of activation by interface mutants. In addition, the interface mutations and PEST domain truncation showed only an additive and not a synergistic increase in basal Notch signal (Figure 4E), further confirming a novel role in Notch signal regulation compared to HD mutants. It is interesting, however, that H1570A differed from H1570P in its inducibility, as the latter was not induced by either extracellular or intracellular stimuli.

**Figure 4.**
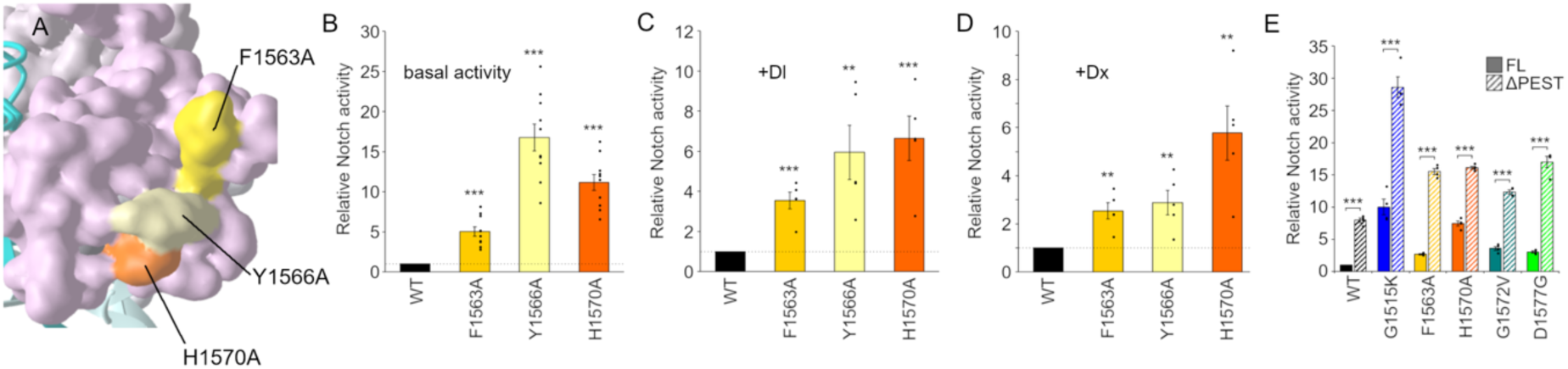
Identification of a novel regulatory hotspot in LNR-C driving ligand-independent Notch activation. **(A)** Human NOTCH1 NRR crystal structure highlighting the positions of three surface-exposed LNR-C mutations (F1563A; amber, Y1566A; pale yellow, and H1570A; orange). **(B)** Strong increase in basal (ligand-independent) Notch signalling induced by the three LNR-C mutations. **(C,D)** Ligand-dependent **(C)** and Dx-induced **(D)** activation of the LNR-C mutants, normalised to wild-type Notch activation under the same conditions. **(E)** Effect of PEST domain truncation on ligand-independent signalling of the LNR-C mutants. P < 0.05 (*), P < 0.01 (**), and P < 0.001 (***) by two tailed t-test, error bars are SEM.

To determine whether the interface mutations affect Notch homodimerisation, we analysed Notch-Notch interaction using a luciferase-coupled co-immunoprecipitation assay (Figure 5A). We indeed detected an interaction between the LNR and full-length Notch, but neither the F1563A nor H1570A mutations showed any reduction in LNR homodimerisation (Figure 5B). This was further confirmed by a split luciferase complementation assay (Figure 5C), where LNR-Nluc and LNR-Cluc demonstrated LNR-LNR interaction, and again, neither F1563A nor H1570A impacted homodimerisation (Figure 5D). Therefore, these interface mutants can strongly elevate multiple Notch signalling pathways without affecting Notch homodimerisation, suggesting that LNR-C mediates a novel regulatory mechanism for Notch signalling.

**Figure 5.**
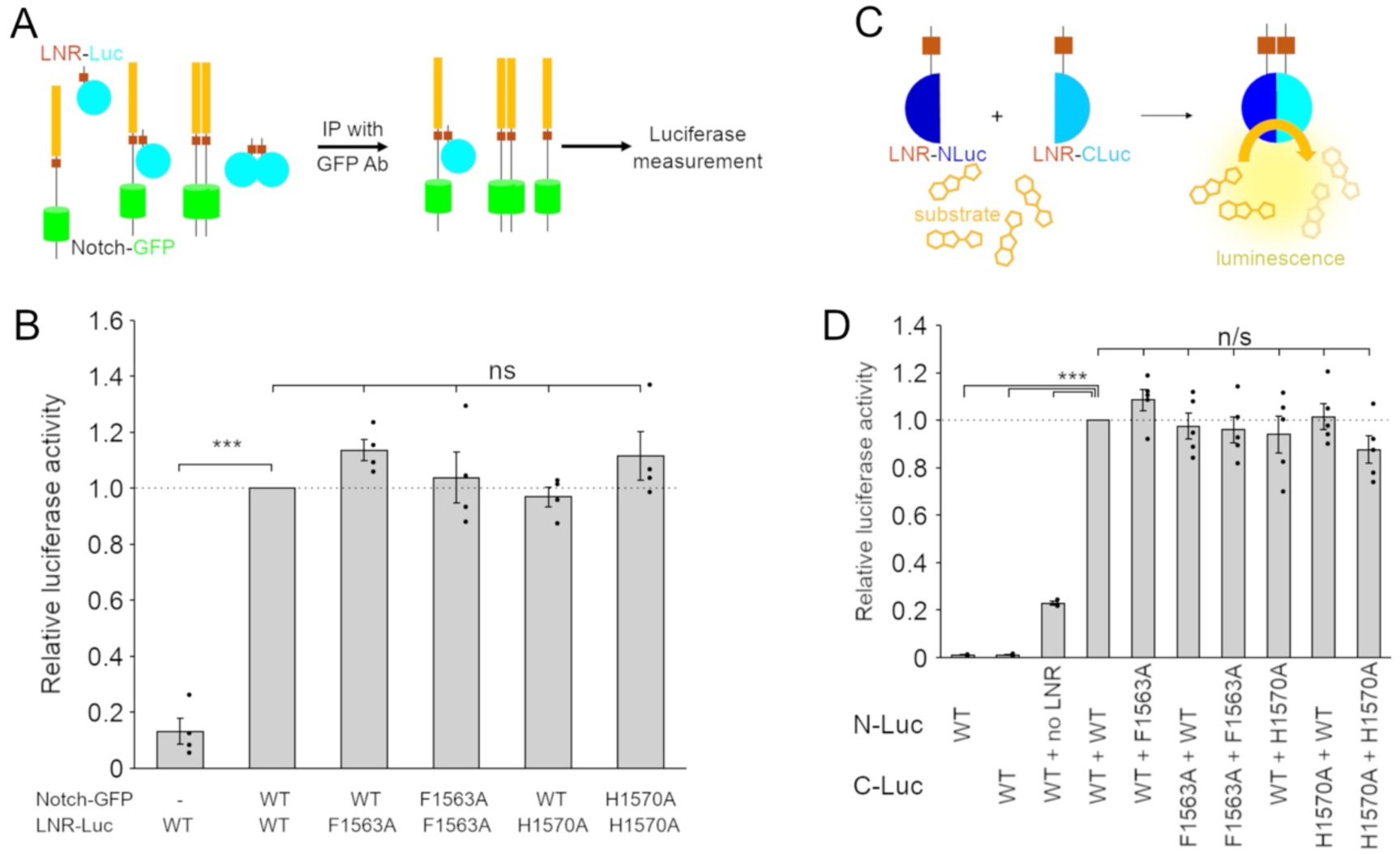
Intact NRR–NRR homodimerisation despite LNR-C interface mutations. **(A,B)** Co-immunoprecipitation–based assay for NRR–NRR homodimerisation. **(A)** Schematic presentation of the assay. EGFP-tagged full-length Notch was immunoprecipitated from S2 cells co-expressing Notch–EGFP and luciferase-tagged Notch LNR domain using an anti-GFP antibody (GFP-trap). Homodimerisation was assessed by measuring co-precipitated luciferase activity. **(B)** Luciferase signals were normalised to total lysate luciferase activity to quantify binding efficiency. Robust NRR–NRR homodimerisation was observed, and none of the tested interface mutations significantly affected this interaction. **(C,D)** Split-luciferase assay to assess the effect of LNR-C mutations on NRR–NRR homodimerisation. **(C)** Schematic representation of the assay. N-terminal and C-terminal fragments of luciferase were fused to Notch LNR constructs and co-expressed in S2 cells. Homodimerisation was detected as reconstituted luciferase activity. **(D)** Firefly luciferase signals were normalised to Renilla luciferase activity to quantify dimerization efficiency. Strong luciferase activity indicated NRR–NRR homodimerisation, and none of the tested LNR-C interface mutations significantly altered this interaction. All constructs include N-terminal signal peptides to ensure proper membrane topology. P < 0.05 (*), P < 0.01 (**), and P < 0.001 (***) by two tailed t-test, error bars are SEM.

To examine whether NRR mutants might affect expression or turnover rate of Notch, the protein expression levels of all the NRR mutants used in this study were examined by western blotting (Figure 6A-D). Except for the E1538R mutant, which exhibited extremely low expression, all the mutants were expressed at levels which were comparable to the wild-type Notch protein, and for the most part we found no significant correlation between basal signal intensity and expression level. However, the LNR Notch homodimerisation interface mutants, H1570A, F1563A, Y1566A, and two *Drosophila* alleles in the same interface G1572V (l1N-B) and D1577G (414) were consistently expressed at a significantly higher level than the wild-type, suggesting that this may contribute to elevated signalling. We note that the H1570P mutation, which did not have Dx or Dl-inducibility also displayed higher Notch protein levels, although not so high as H1570A. These observations indicate that altered stability or degradation is not a universal mechanism underlying the diverse signalling outputs of NRR mutants, but rather a feature restricted to this specific structural subset. For most mutants, changes in signalling occur independently of protein abundance, consistent with the idea that they act primarily by modifying NRR autoinhibition, conformational dynamics, or ADAM-family metalloprotease accessibility rather than by affecting global turnover. Next, we measured the degradation kinetics of the Notch mutant proteins using a cycloheximide chase assay (Figure 6E,F). Degradation of the wild-type Notch protein began within 4-6 hours of cycloheximide addition, whereas both F1563A and H1570A displayed a significantly slower degradation rate. In contrast, activating mutants G1515K and E1705P that are located in the LNR/HD interface region, and the ΔPEST construct, showed similar degradation kinetics to WT (Supplementary Figure S3). This suggests that the LNR-C interface is a novel functional domain involved in regulating Notch protein stability and degradation with an impact on Notch activation, likely contributing, at least in part, to the increased intensity of both basal and induced Notch signals.

**Figure 6.**
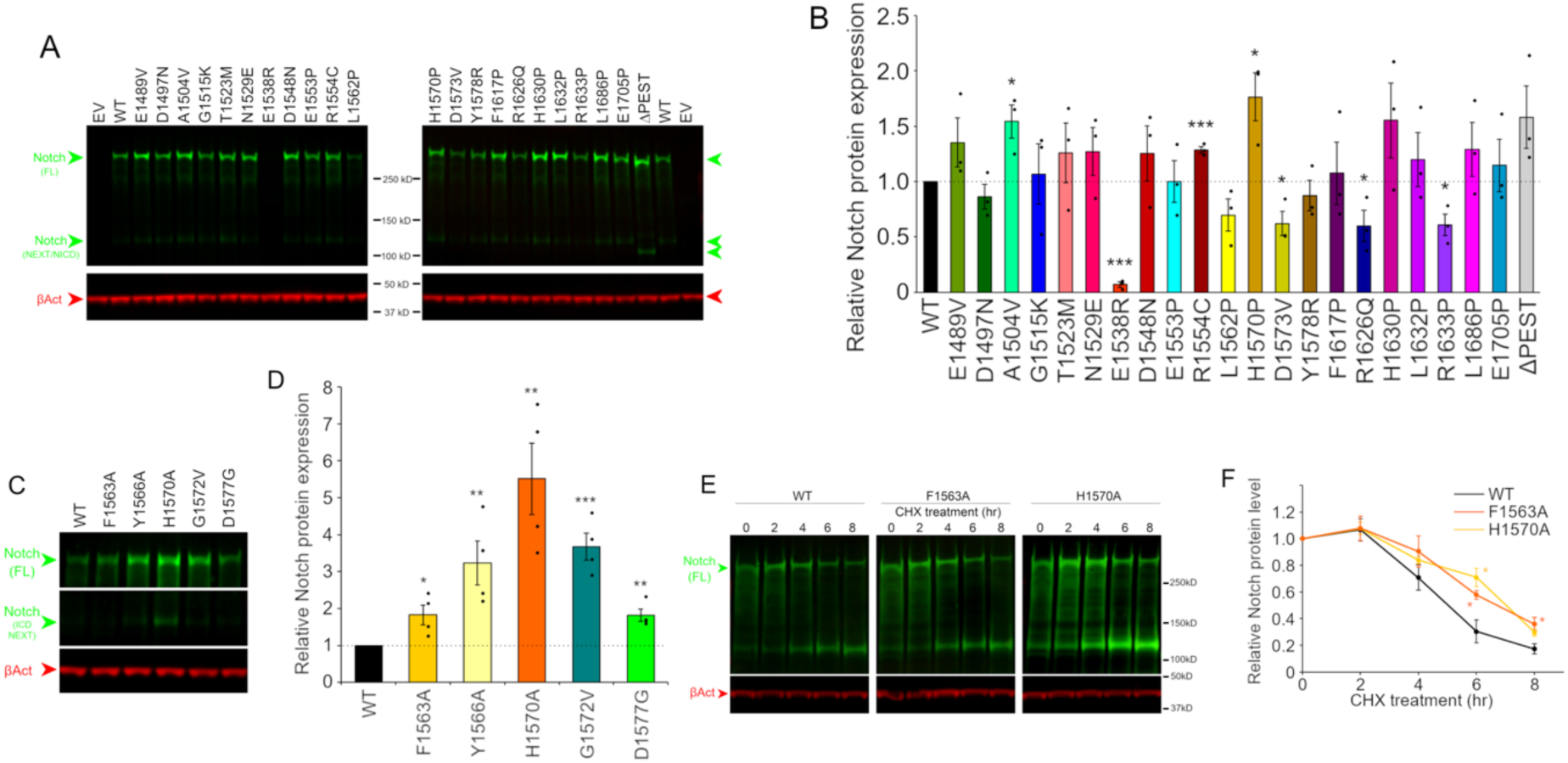
A novel negative regulatory function of LNR-C in Notch protein stability. **(A,B)** Protein expression levels of Notch cancer mutants in S2 cells. Expression profiles of all cancer-derived Notch mutants used in this study were examined by western blotting **(A)** and the band intensities of full-length Notch were quantified to compare relative protein levels **(B)**. **(C,D)** Elevated Notch Protein Levels by LNR-C Interface Mutations. Expression profiles of LNR-C interface mutants and two Drosophila alleles, l1N-B (G1572V) and 414 (D1577G), were examined by western blotting **(C)** and the band intensities of full-length Notch were quantified to compare relative protein levels **(D)**. **(E,F)** Enhanced stability of Notch LNR-C interface mutants revealed by cycloheximide (CHX) chase assay. **(E)** S2 cells expressing LNR-C interface mutants (F1563A or H1570A) were treated with 10 µM cycloheximide for the indicated time and assessed by western blotting. **(F)** Band intensities of full-length Notch were quantified to evaluate protein stability over time. P < 0.05 (*), P < 0.01 (**), and P < 0.001 (***) by two tailed t-test, error bars are SEM.

## Discussion

The NRR of Notch plays a pivotal role in the Notch activation mechanism by sequestering the S2 cleavage site in the absence of signal induction which keeps basal activation levels low (Gordon et al., 2009; Gordon et al., 2007). In humans many Notch mutations in the NRR have been reported in both solid tumours (Aster et al., 2017; Mutvei et al., 2015; Zhang et al., 2016) and the blood cell cancers (Rosati et al., 2018; South et al., 2012), such as T-ALL (Belver and Ferrando, 2016; Pear and Aster, 2004). Despite the important role that *Drosophila melanogaster* has played in probing Notch structure/function, there have been few studies of NRR mutations (Brennan et al., 1997; Lyman and Young, 1993). Here we generated several mutations located in different regions of the NRR of *Drosophila* Notch to investigate their consequences on Notch activation and compare their behaviour to previously studied T-ALL mutations. We generated mutations in the core HD, that were similar to those most commonly found in T-ALL, mutations in the LNR/HD interface and mutations in the LNR domains. The latter included perturbation of a putative dimerisation interface that was previously identified through crystallographic studies of human Notch (Gordon et al., 2009; Xu et al., 2015). It is intriguing that no Notch alleles equivalent to T-ALL mutations in the HD domain have been reported in the fly. This stands out especially when considering the existence of multiple lines of gain-of-function alleles, such as *Abruptex* (Ax, missense mutations in EGF24-29) (Foster, 1975; Portin, 1975) and *Microchaetae defective* (*Mcd)*, nonsense mutations in the C-terminal region including in the PEST (Ramain et al., 2001). In this study, we demonstrated that, unlike in human NOTCH1, certain HD mutations, when introduced into *Drosophila* Notch, accumulate around ER/Golgi and do not increase ligand-independent basal Notch activity and are not induced by Dx or Dl. This suggests that this class of mutation destabilises the HD fold, triggering retention in the secretory pathway, making it unavailable for signal activation. The reason for the difference with human Notch is still unclear but could be related to the absence of S1 cleavage (Kidd and Lieber, 2002) in *Drosophila* Notch since this class of mutation is thought to destabilise the HD-N/HD-C interaction (Malecki et al., 2006). It is possible that Human NOTCH1 L1600P and L1678P variants remain surface-expressed because S1 cleavage generates a pre-assembled HD-N/HD-C heterodimer that masks destabilised regions. In contrast, because *Drosophila* Notch folds and traffics as a single polypeptide, introduction of analogous HD-destabilising substitutions (L1632P and L1686P) may expose hydrophobic residues that impair maturation, resulting in ER/Golgi retention. However, currently our results are restricted to S2 cells and it is possible that *in vivo*, in some cell contexts, then cells are more tolerant and allow some transport to the cell surface and activation. Future work will address this possibility.

Interestingly, mutations affecting residues that lie in the interface between the LNR and HD domains, R1626Q and E1705P, showed elevated basal signalling compared to WT. Furthermore, when combined with PEST domain truncation, these mutations produced a synergistic increase in activity, indicating their similarity in behaviour to T-ALL mutations. These mutations showed little or no inducibility in signal activity either by extracellular ligand or by intracellular interaction with Dx. The most likely explanation is that these mutations already caused full exposure of the S2 site for metalloprotease cleavage. Additionally, we tested four *Drosophila* NRR domain mutants and found similarly elevated constitutive signalling. These included three mutants previously shown to be gain of function *in vivo*. l1N-B (G1572V) (Lyman and Young, 1993) and 414 (D1577G) (Brennan et al., 1997) are classical *Drosophila* Notch alleles, the latter also being a NOTCH1 T-ALL mutation. G1572, located in LNR-C, interacts with the HD domain, and its mutation likely disrupts the LNR-HD interaction. D1577 is also located in LNR-C and its mutation most probably disrupts calcium binding (Gordon et al., 2009) involved in stabilising the LNR structure. The CC-SS mutant (Lieber et al., 1993) removes a conserved disulphide bond to destabilise the HD and has previously shown to activate Notch signalling when overexpressed in fly tissue. In addition, a triple mutant LGI-AAA (Gordon et al., 2007), which removes a cap over the S2 site allowing constitutive access to ADAM10-dependent proteolytic cleavage, resulted in strongly elevated basal Notch signalling. As with the HD/LNR-C T-ALL mutants R1626Q and E1705P, none of the four *Drosophila* mutants were significantly inducible by Dx of Dl. There was a context dependence regarding the impact of different mutants in Notch activity compared to WT. While all the above mutants stimulated constitutive basal activity, in the presence of Dx or Dl there were different outcomes for different mutants. In the presence of Dx or trans-presented ligand Dl, R1626Q and E1705P had reduced signalling compared to WT in equivalent conditions, whereas the CC-SS and LGI-AAA remained at least as active as WT Notch stimulated by Dx or Dl, because of their higher relative basal signal. Interestingly, while L1N-B and 414 also retained higher activity than WT in the presence of Dl (significant for L1NB), in the presence of Dx the activities of both mutants were significantly reduced compared to WT, indeed Dx caused a significant reduction in activity of 414 compared to its own basal activity. Dx has been shown to promote Notch endocytosis and to divert Notch into a clathrin-rich endosomal membrane subdomain (Shimizu et al., 2024) where it may not be accessible to ADAM10 and perhaps this may explain the partial suppression of Notch activation in this case. If a similar context dependence translates to human Notch cancer mutants, then this may suggest ways to modify oncogenic properties in a mutant-specific manner, and it would be interesting to explore this possibility further. It is not clear, however, why the impact of Dx to down regulate basal Notch activity is specific to the 414 allele. It would be interesting to determine whether Dx has different outcomes on the endocytic trafficking of Notch for different mutants.

In addition to HD mutants, we investigated the consequences of mutations affecting the LNR domains A-C. A range of different responses were identified producing both increased and decreased basal signal activation and variable degrees of inducibility in response to Dl and Dx. G1515K, located in LNR-A adjacent to the S2 site, and E1553P, located in LNR-B adjacent to the HD disulphide bond, behave similarly to activating HD mutants and to the *Drosophila* alleles, i.e. they displayed strong constitutive basal signalling, no inducibility with Dl or Dx, and synergistic activation after removal of the PEST region. Due to the very high basal signalling these mutants remain significantly more active than when WT is induced by Dl or Dx. Other LNR mutants located on the domain surface but on the opposite side to the HD interface had more subtle affects. For example E1489V, located in LNR-A shows only a small increase in basal activation but its signalling levels are higher compared to WT in Dl or Dx-inducing conditions, whereas E1538R behaves as loss of function in all conditions, both basal and induced, and shows no inducibility in response to Dx or Dl. D1548N, located in LNR-A, shows normal basal activation, but has decreased activity in inducing conditions due to lack of inducibility. However, D1573V, located in LNR-C, displayed higher basal activation but in inducing conditions activity was significantly less than WT, also due to absence of inducibility.

A particularly intriguing region of LNR-C is a set of conserved residues that occupy a region designated from crystal structures as a putative Notch/Notch dimerisation site. Mutations of this region in human NOTCH3 increased ligand-independent basal Notch activation (Xu et al., 2015). We mutated conserved residues in this interface in *Drosophila* Notch and also found significantly increased basal signalling. Despite strong basal activation of two of these mutants, F1563A and H1570A were similarly inducible as WT by both Dx and Dl and their overall activity in these conditions was substantially higher than induced WT signalling, in contrast to activating HD mutants which were not further induced from a high basal level. Another difference was that although mutation of the PEST domain increased the activity of these LNR-C interface mutants this was additive and not synergistic. These results suggest a different mechanism of signal upregulation than simple exposure of the S2 cleavage site. We found that the Drosophila LNR domain was sufficient to mediate dimerisation, but this was not perturbed by the interface mutants. Instead, we found that these mutations upregulated Notch protein levels and had decreased degradation rates compared to WT. This suggests the existence of a regulatory mechanism that facilitates Notch protein degradation through this LNR-C interface region, providing broader control of Notch signalling, although we don’t rule out additional regulatory interactions of this interface. Understanding how the LNR-C interface contributes to Notch downregulation could open a new strategy for therapeutic modulation of Notch activity, offering potential treatments for diseases driven by aberrant Notch signalling, such as T-ALL. Both H1570A and H1570P substitutions increased Notch protein levels, which is reflected in their elevated basal signalling. H1570A shows the highest basal signal, consistent with its stronger stabilization, whereas H1570P shows only a moderate increase. The reduced activation of H1570P, despite its stabilization, may reflect a negative structural effect of the proline substitution that interferes with NRR conformational changes, leading to impaired ligand- and Dx-induced signalling. These observations suggest that, in addition to stabilizing the protein, the nature of the substitution can influence the receptor’s responsiveness to activating cues.

Our results therefore illustrate that there is considerable variety of outcomes of NRR cancer mutants on the mechanisms of Notch signalling and the levels of signal output in different cell contexts compared to WT. This mechanistic variety must therefore be considered when developing optimum treatment strategies. As these experiments were performed in S2 cells, which provide a simplified but non-hematopoietic environment, the system is best viewed as an initial platform to compare the intrinsic regulatory properties of different NRR mutations before moving to more physiological contexts, such as the *Drosophila* hematopoietic system *in vivo*. However, S2 cells lack some cofactors, including Fringe, specific DSL ligands, and specialised cell junctions, that influence Notch activation. Thus, while conserved features such as NRR autoinhibition and ligand-induced conformational control can be reliably examined in this system, the quantitative effects of individual mutations on signal strength or sensitivity will need to be evaluated in more physiological settings. This can be done by either expressing activating mutant cDNA constructs, or gene editing mutants into the endogenous *Drosophila* Notch gene, although the latter may cause a lethal phenotype. Despite conserved mechanisms of both Notch signalling and the hematopoietic stem cell system in humans and *Drosophila* (Banerjee et al., 2019; Boulet et al., 2018; Gold and Bruckner, 2014), a model system to study T-ALL has not been established in *Drosophila*. Although the most common HD mutations in *Drosophila* Notch appear not to mimic Notch activation in T-ALL, our study identifies subsets of mutation that could be used to model T-ALL in *Drosophila* which could complement existing mammalian models to significantly advance our understanding of the disease and accelerate therapeutic discoveries. In particular, a subset of mutations was found which recapitulates the synergism observed between NRR and PEST region mutants that is a notable feature of human T-ALL. Establishing an *in vivo* model would enable high-throughput genetic screens to identify novel genes and pathways, provide a platform for rapid drug target validation and even for investigation of treatment resistance mechanisms. Understanding different mechanisms of misregulation that are specific to different classes of mutation will enable development of personalised treatment options.

## Materials and methods

### Cell culture

Drosophila S2 cells were obtained from Thermo Fisher Scientific (Waltham, MA, USA) and were cultured in Schneider’s *Drosophila* medium (Sigma-Aldrich, St. Louis, MO, USA) supplemented with 10% heat-inactivated fetal bovine serum (Thermo Fisher Scientific) and 0.5% Penicillin-Streptomycin (Merck) at 25°C. Effectene (Qiagen, Manchester, UK) was used for transfection.

### DNA constructs

The pMT–Notch–EGFP and pMT–Dx–V5 constructs were previously described (Shimizu et al., 2024). All point mutations and truncations were introduced into the pMT–Notch–EGFP backbone. For the pMT– LNR–Luciferase construct, a bovine prolactin signal peptide was fused to a Drosophila Notch LNR domain fragment (residues G1465 to V1600), followed by a flexible linker (GGGSSGG), the firefly luciferase open reading frame, and a stop codon. For the split-luciferase assay, the N- and C-terminal fragments of firefly luciferase (E2–G416 and S399–K547, respectively) were fused to the pMT–LNR construct to generate pMT–LNR–NLuc and pMT–LNR–CLuc. PEST domain truncation was achieved by removing the C-terminal region of the Notch–EGFP construct through StuI restriction enzyme digestion.

### Luciferase Assay

*Drosophila* S2 cells (∼1 × 10⁶ cells/ml) were transfected using Effectene (Qiagen) according to the manufacturer’s instructions with 1 ng of Notch construct, 500 ng of NRE-firefly luciferase reporter, and pMT-renilla luciferase plasmid as a transfection control (Shimizu et al., 2024). After 48 hours, cells were transferred to a Nunc 96-well assay plate (Thermo Fisher Scientific), and Notch expression was induced by adding 1 mM CuSO₄ for 24 hours. Luciferase activity was measured using the Dual-Glo Luciferase Assay System ((Promega, Southampton, UK) on a luminometer (Berthold, Bad Wildbad, Germany). Firefly luciferase activity was normalised to renilla luciferase activity for all analyses. For Dx-mediated Notch activation, 10 ng of Dx-expressing construct was co-transfected with the reporter system. For ligand-induced Notch activation, Dl expression in S2-Dl cells was induced with 1 mM CuSO₄ for 3 hours in a 96-well plate. Cells were then fixed with 4% formaldehyde (Polyscience, Warrington, PA, USA), quenched with 100 mM glycine, and co-cultured with Notch-expressing cells in the presence of 1 mM CuSO₄ for 24 hours. For the split-luciferase assay, 500 ng each of LNR–NLuc and LNR–CLuc constructs were co-transfected into S2 cells along with a renilla luciferase plasmid as a normalization control.

### Immunofluorescent microscopy

Transfected S2 cells were seeded onto poly-L-lysine–coated coverslips and fixed with 4% formaldehyde (Polyscience) in PBS. Cells were permeabilised using 0.2% Triton X-100 (Sigma-Aldrich) in PBS, followed by blocking in 3% skim milk in PBS. Cells were incubated with primary antibodies: mouse anti-PDI (Kondylis et al., 2001) (1:500; monoclonal clone 1D3, Enzo Life Sciences, Farmingdale, NY, USA) and rabbit anti-GM130 (Kondylis et al., 2001) (1:500; polyclonal, Abcam, Cambridge, UK). Secondary antibodies used were Alexa Fluor 568–conjugated anti-mouse IgG and Alexa Fluor 647– conjugated anti-rabbit IgG (Jackson Immuno Research Ely, UK).

### Co-immunoprecipitation

S2 cells were transfected with pMT–Notch–EGFP and pMT–LNR–Luciferase constructs in a 6-well plate. After expression, cells were lysed in 150 µL of lysis buffer (100 mM Tris-HCl (pH 7.5), 150 mM NaCl, 1% Tergitol 15-S-9, 1 mM CaCl₂, 50 µM MG132, and Halt Protease and Phosphatase Inhibitor Cocktail, Thermo Fisher Scientific)). Lysates were incubated with GFP-Trap beads (ChromoTek, Planegg, Germany) for 1 hour at 4 °C to immunoprecipitate Notch–EGFP. Luciferase activity in the co-immunoprecipitated LNR–Luciferase, as well as in 10% input lysates, was measured using a luminometer (Berthold).

### Cycloheximide (CHX) chase assay

S2 cells expressing Notch–EGFP were treated with 10 µM cycloheximide (Sigma-Aldrich) in culture medium lacking CuSO₄ for the indicated time periods. Cells were then lysed using the lysis buffer. Notch protein levels were assessed by western blotting using a mouse anti-Notch intracellular domain (ICD) antibody (1:10,000; monoclonal clone C17.9C6 (Fehon et al., 1990), Developmental Studies Hybridoma Bank, University of Iowa, IA, USA) and a rabbit anti–β-actin antibody (1:10,000; polyclonal, Proteintech, Manchester, UK) as a loading control. Detection was performed with the LI-COR Odyssey imaging system (LiCor Biosciences Lincoln, NE, USA) using Alexa Fluor Plus 800– conjugated anti-mouse IgG and Alexa Fluor Plus 680–conjugated anti-rabbit IgG secondary antibodies (1:10,000; Thermo Fisher Scientific).

### Statistical analysis

Data are presented as mean ± SEM. Statistical significance was assessed using a two-tailed Student’s t-test. Significance levels are indicated as P < 0.05 (*), P < 0.01 (**), and P < 0.001 (***). Data were assumed to follow a normal distribution; however, this was not formally tested.

## Acknowledgments

We acknowledge financial support from the BBSRC, grant no. BB/V014218/1 and thank Sarah Bray, University of Cambridge, UK for providing the NRE-luciferase plasmid and the Developmental Studies Hybridoma Bank, University of Iowa, for antibodies.

