## Supplementary figures S1-S3 for "Analysis of cancer mutations introduced into the Drosophila Notch Negative Regulatory Region uncovers a diversity of regulatory outcomes"

**Supplementary information**

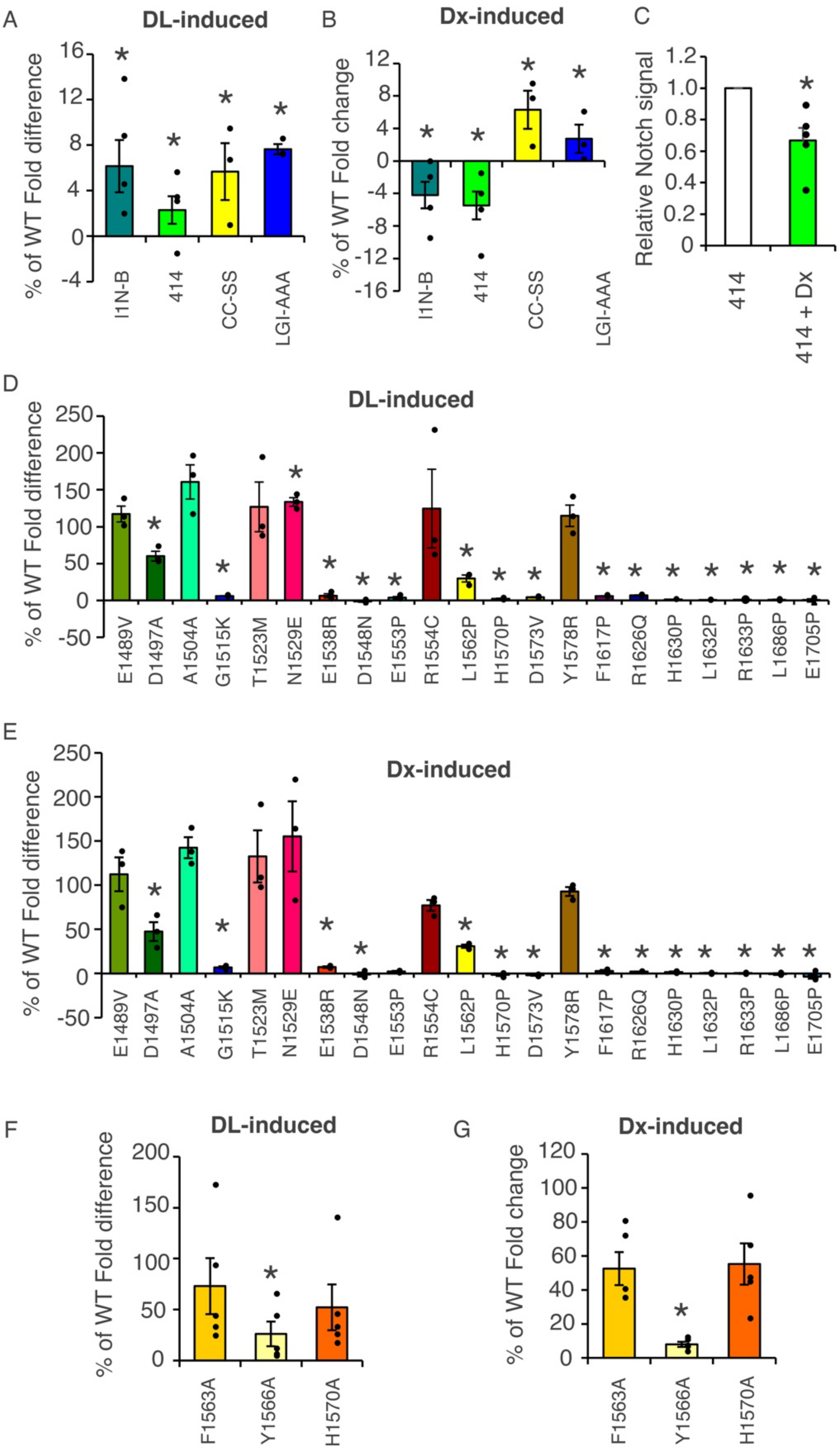


**Supplementary figure S1: Inducibility of NRR mutants by Dl and Dx compared to WT. A,B,** Inducibility of NRR mutants compared to WT for Dl-induced **(A,D,F)** and Dx-induced **(B, E,G)**

expressed as a % of WT. **(A-C)** Inducibility of *Drosophila* NRR mutant alleles to Dl **(A)** and Dx **(B)**. **(C)** Fold-decrease of 414 allele after Dx expression compared to 414 basal shows significant decrease. **(D,E)** Inducibility of cancer-associated HD and LNR domain mutants for Dl **(D)** and Dx induction. **(F,G)** Inducibility of LNR-C dimer interface mutants in response to Dl **(F)**, and Dx **(G)**. To calculate relative inducibility first all basal signals were normalised to 1. The "fold difference" was calculated as induced signal -1 for mutant and for WT and mutant value expressed as a percentage of the WT fold difference. Hence 100% indicates fold inducibility of mutant is same as WT, 0% indicates mutant signal does not change in induced conditions compared to its own basal signal, and negative numbers are cases where the mutant signal is reduced in the presence of Dl or Dx, compared its own basal condition with the fold difference expressed as a % of the WT fold difference. * indicates $P<0.05$ compared to WT **(A,B,D-G)** or compared to 414 basal **(C)**, by 2 tailed t-test, error bars are SEM.

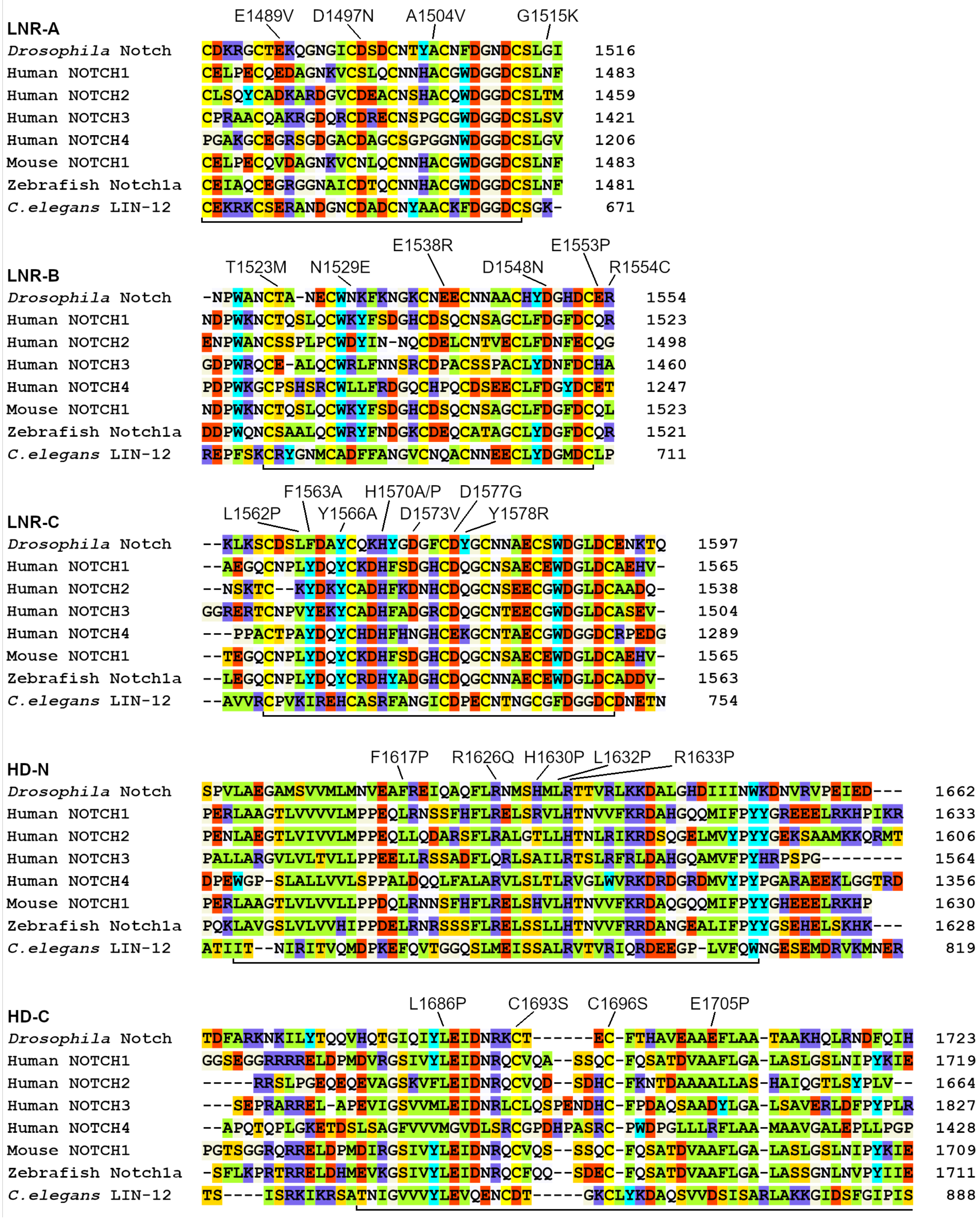


**Supplementary figure S2: Positions of NRR point mutations used in this study, shown in the context of amino-acid conservation.** Alignment of the five domains (LNR-A/B/C and HD-N/C) from different Notch proteins (*Drosophila* Notch, human NOTCH1–4, mouse NOTCH1, zebrafish Notch1a, and *C. elegans* LIN-12) was colour-coded: green for hydrophobic residues, red for acidic, violet for basic, yellow for cysteine, orange for serine and threonine, cyan for tyrosine and tryptophan, light purple for asparagine and glutamine, and light yellow for glycine and proline.

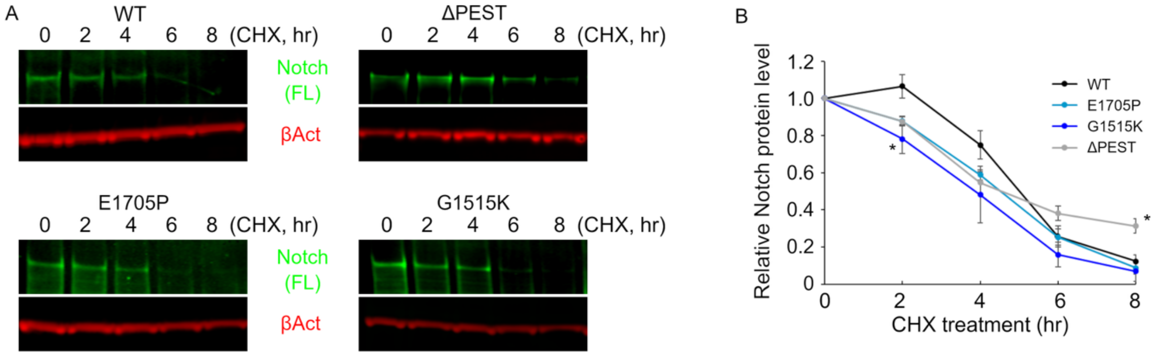


**Supplementary figure S3: LNR–HD interaction mutations do not affect Notch protein stability, as revealed by a cycloheximide (CHX) chase assay.** (A) S2 cells expressing LNR–HD interaction mutants (G1515K and E1705P) or a PEST domain–truncated variant were treated with 10 μM cycloheximide for the indicated times and analysed by western blotting. (B) Band intensities of full-length Notch were quantified to assess protein stability over time. * indicates $p<0.05$ compared to WT (2 tailed t-test), error bars are SEM.
